# Recent developments in the production of 2D- and 3D colon and stomach adenocarcinomas primary cell models

**DOI:** 10.1101/2023.02.26.529716

**Authors:** Igor Tokarchuk, Oleksandr Mushii, Alona Dreus, Anastasia A. Koziaruk, Dmyto O. Shapochka, Oleg A. Ryzhak, Serhii O. Venhryk, Iurii S. Golovko, Sergey V. Ryabuhin, Anton I. Hanopolskyi, Duncan B. Judd, Dmitriy M. Volochnyuk

**Affiliations:** Preci LLC, 9 Amosova Street, 03038 Kyiv, Ukraine; Feofaniya Clinical Hospital, 21 Akademika Zabolotnoho Street, 03143 Kyiv, Ukraine; Biopharma Plasma Ltd., 6 Amurska Street, 03022 Kyiv, Ukraine; Kyiv Regional Oncology Center, 1 Bahhovutivska Street, 02000 Kyiv, Ukraine; Enamine Ltd., 78 Chervonotkatska Street, 02660 Kyiv, Ukraine; Institute of High Technologies, Taras Shevchenko National University of Kyiv, 60 Volodymyrska Street, 01033 Kyiv, Ukraine; Institute of Organic Chemistry, National Academy of Sciences of Ukraine, 5 Murmanska Street, 02660 Kyiv, Ukraine; Awridian Ltd., Stevenage Bioscience Catalyst Gunnelswood Road, Stevenage Herts, SG12FX, UK

**Author notes:** (I.T.); (O.M.); (A.D.); (O.A.K.); (A.I.H.). (D.O.S.). (O.A.R.). (S.O.V.); (I.S.G.). (S.V.R.); (D.M.V.). (D.B.J.).

**Keywords:** primary cell cultures, patient-derived cells, gastric cancer, colorectal cancer, organoids, spheroids

## Abstract

Gastric and colorectal cancer models are essential for the advancement of precision medicine discovery and development. 2D attached monolayer, spheroid and organoid approaches have all been used in the formation of biobanks containing primary patient-derived cells. Here, we report an assessment of those procedures for a panel of nine patient-derived adenocarcinoma samples, along with the most applicable method for the bio-banking of these cell types. A live cell biobank of tumour specimens would facilitate drug discovery laboratories to evaluate drugs on the population of cell cultures, prior to the clinical phase.

## 1. Introduction

Cancer research utilizes a wide range of disease models (1), but only a few accurately reflect human disease (2). For example, immortalized cancer cell lines used in research rely on abnormal cell cycles, which results in the accumulation of genetic mutations. Furthermore, the need for artificial attachment and culture media alters the phenotype of those cells (3,4). Despite their limitations, the technical simplicity of these models has enabled them to remain a popular choice (5,6). In contrast, primary cell models are technically more challenging but offer the potential to reflect human disease better (7). However, both technical feasibility and the representation of human diseases are difficult to achieve simultaneously. It is recognized that low-passage cancer cells derived from patients are a valuable source of knowledge about cell physiology, as well as cell-cell and cell-matrix interactions (8). Among them, 2D attached monolayer cells (9) are the simplest but are challenging to apply to epithelial-like cancer cell models. Thus, 3D spheroids (10) were developed as artificially crowded cell conglomerates, and these display the spatial distribution of cell types and recognizable physiological features., In addition, 3D organoids (11), which self-assemble upon differentiation of stem cells, also reresent complex morphology. However, these are more expensive to generate and characterize (12,13). All of these models can be defined as primary if derived directly from living organisms. The inability to scale up these models because of low proliferation rates and cellular senescence limits their wider use (14). Cell expansion limitations also frequently restrict experiment cross-validation and reproducibility assessment. More recently developed organoid models faithfully recapitulate the tumour morphology and can be expanded and used in drug response assays (15).

Spheroid 3D culture is a known source of knowledge of drug effect on the tumour phenotype in a morphological sense (16). Spheroid Cultures can incorporate not only tumour cells, but also immune or stromal cells. In this fashion key importance is relied upon Cancer-Associated Fibroblasts (17–20). Obviously, 3D spheroids can be assembled from immortalized cell lines. However, if viable primary cell lines can also be assembled within spheroids, with obviously larger technical challenges (21). Isolation of cancer-associated fibroblasts (CAFs) is an active task for creation of tissue mimicikng systems in oncology. Immortalized CAFs have been used in numerous works (22–24), however, the issue of faithfulness of those is outstanding. Here we also report the procedures for tissue-derived CAFs explantation and cultivation. Also, co-cultures with cancer spheroids were attempted.

Our initial focus was to evaluate procedures that could enable the expansion of primary gastrointestinal adenocarcinoma cells (GIAC) and, if successful, could be applied to create a living biobank: specifically a collection of assay-ready, proliferating cells for drug evaluation or target identification on a population of genetically unique models (25). The value of using cell line panels (albeit with immortalized cell lines) is illustrated with the MDM2/MDMX inhibitor ALRN-6924. In this study, the inhibitor appeared effective against the majority of wt-TP53 lines. However, it lacked activity against many of the mutated or null TP53 cell lines (26). Thus, such cell line-based screening might be a valuable tool for pharmacogenomic evaluation of leads during the development stages (27).

Over a quarter of cancer cases are gastrointestinal tumours, of which more than a third account for cancer deaths (28,29). Hence the interest in organ modeling for drug discovery and biomarker research. Of particular note are recent advances in organoid production resulting in realistic *in vitro* modeling of colorectal cancer with a heterogeneous cell-based profile (30).

Although 2D monolayer cancer-derived cells are often used in imaging assays (31), there are conflicting reports of their limited expansion potential. Jeppesen et al. conducted a comparative study of colorectal cancer cell extraction and cultivation procedures as monolayer cultures (32) and concluded that trypsin was the best method to digest the tissue. However, short-term cultivation limited by the low expansion potential of cellular replication may result in insufficient numbers of cells for a robust assay. Large multi-cancer research study also showed the potential of tissue culture expansion using foreskin fibroblasts. Among all of the tissues mentioned in that study, only a quarter of cell cultures were successfully expanded (33).

Considering their potential, we investigated 3D organoid models as well. There are many examples of the successful expansion of colorectal and stomach cancer organoids using specialized media and attachment conditions. Short-term cultures of tumour-derived organoids have already proven their effectiveness in personalized medicine over a wide range of tissues (34). Although there is a lack of reported live-cell biobanking of 3D spheroids, they are conceptually applicable in this field as well. Miyoshi et al. showed the expansion of more than 100 patient-derived spheroids with the use of specific media (35).

Despite the overwhelming number of approaches, the choice of method for assay on primary cells still causes confusion from the reproducibility, accessibility and other practical issues. The general aim of this paper is comprehensive comparison of existing methodologies, using patient-derived cancer cells. This work was a part of our internal project of living cell biobanking of cancer cells. This problem was critical for us in the course of creation of reproducible homogeneous Standard Operational Procedure for cancer cell banking. We decided to publish current results, taking into account the generality of this problem for the worldwide community. Here we report an applicable procedure for upscaling and subsequent preservation of GIAC models. We have applied a success-rate-based approach to evaluate the scalability potential of nine patient-derived cell populations. This work also covers the expansion potential of cancer-associated fibroblasts (CAFs) from GIAC. Current work intentionally omits the issue of representation of the actual disease. However, we will try to analyse this aspect as well.

## 2. Materials and Methods

### 2.1. Human Biological Samples

Tumour specimens were provided by Kyiv Region Oncological Center, following the Local Ethical Committee’s approval (January 2021). Tumours were resected according to the standard protocols, immediately washed with 0.9% NaCl solution in the surgery room, suspended in the Transport Media, cooled to 4°C and delivered to the laboratory within 2 hours post resection.

### 2.2. Transport Media

Custodiol (Dr. Franz Köhler Chemie GmbH) – 25 mL per sample

Normocur (Invitrogen) – 100 μg/mL

Primocin (Invitrogen) - 100 μg/mL

Fungin (Invitrogen) - 50 μg/mL

### 2.3. Pathological FFPE block preparation

After receipt of samples at the laboratory a 3 mm piece of tissue was submerged in Formalin (10% Formaldehyde), subsequently embedded in paraffin using the classical method, washed with Chloroform to remove Formalin, followed by Ethanol. The sample was submerged in Paraffin-Xylene (1:1 ratio) solution, and this solution was replaced with pure liquid thawed paraffin (55°C) and let to solidify at room temperature. Blocks can be stored indefinitely for at least 5 years for protein and DNA analysis.

### 2.4. Histological Analysis

For histological staining: the Hematoxylin-Eosin method was used (36). The pathohistological analysis was performed using light microscopy. The latest WHO (37) classification was used to define cancerous tissue, describing its TNM and Grade.

### 2.5. Tissue Processing and Digestion

All necrotic regions, blood vessels, and fat were resected, and the sample was washed with Hanks buffer. For cancer cell extraction the epithelial layer was excised and then washed. For CAF extraction the submucosal layer was retained.

The whole sample was washed with the Washing Solution, which consists of DMEM (BioWest) with antibiotics and antimycotics (Normocur 100 μg/mL Primocin 100 μg/mL, and Fungin 50 μg/mL, all from Invitrogen). Then it was additionally washed with DPBS.Before digestion, tissue was minced with surgical scissors to the size of 1-2 mm. Digestion was performed in a mixture of Collagenase IV (STEMCELL Technologies) + Hyaluronidase (Sigma) or Dispase II (Sigma) solution. Digestion was performed from 1 to 3 hours depending on sample viscosity with occasional mixing.

Cell numbers were determined using a counting chamber, with Trypan Blue (Sigma) viability stain.

Following digestion, the cell suspension was filtered through a cell strainer (either 40 or 70 μm), washed three times with DPBS and seeded into pre-coated plates.

Culture media (per 10 ml) consists of:

8,960 ml DMEM (BioWest) with 0.5 mL FBS (Gibco) - 5%

EGF (Invitrogen) – 20 μg/mL,

B-27 (Gibco) - 1x, 200 μL of stock,

L-Glutamine (Gibco) - 2mM, 100 μl of stock,

HEPES (Sigma) - 2mMInsulin (Sigma) – 7.5 μg/ml, 38 μL of stock,

Pen/Strep (Gibco) - 1x, 100 μL of stock.

### 2.6. Well-Plate Coating with BME

For the well surface coating: BME (Cultrex) aliquots were thawed on ice. DMEM, plates, and tips were pre-cooled to 4°C. A coating solution of 1 part BME and 40 parts DMEM (by volume) was prepared. Using precooled tips, 150 μL of coating solution was dropped into each well and immediately removed. This was repeated three times. One portion of the coating solution (150 μl) was reused for up to three times and then discarded. The coating solution temperature was maintained at less than 4°C. The well-plate was placed into the incubator (37°C) for 10 minutes. Each well was visually inspected to ensure a coating of BME was visible. Plates were stored at 4°C and used within 7 days.

### 2.7. Cultivation Procedure

For 2D cell culturing the media was replaced every 3 days. After achieving a local confluent monolayer, passage was performed and cells were usually expanded to a density of around 50%.

### 2.8. Fibroblasts Isolation Procedure

For fibroblast isolation, explant technique was used. After the washing step (Washing Media is described in 2.5), tissue was digested in Dispase II for 30 minutes. Next, the tissue is cut into small (2-3mm) pieces. The tissue pieces were suspended in a plate or T25 flask containing Culture Media. After 3-4 days all migrated cells were detached, and the suspension was washed through a cell strainer. The resulting cell suspension was seeded into pre-coated wells.On visual detection of fibroblasts in the cell population, TrypLE Express (Thermo Fisher Scientific) was used for differential cell detaching. TrypLE Express was added to the well for 5 min. Detached epithelial cells were discarded to the separate vial. The remaining cells were predominately fibroblasts which had been detached from the surface for 10-15 minutes.

### 2.9. 3D spheroid cultivation procedure (ULA)

After the digestion step, cells were seeded into ULA plates (CellCarrier Spheroid ULA 96-well Microplates) (100k cells per well) with 50% of media volume exchanged every second day. Cells were passaged after spheroid size on the bottom grew twofold.

### 2.10. Cell viability analysis

Cell viability was analysed using Trypan Blue staining with 0.5% solution in water (1:1 with cell suspension). Ethanol-fixed cells were used as a positive control. QC for the plated cells was done with 10 mg/mL solution of Propidium Iodide in water via addition of 2 μL of this solution to the 100 mL well.

## 3. Results

### 3.1. GIAC tissue collection and processing

Fresh tissues were procured under Local Ethical Committees approval and following Ukrainian GCP requirements (see Materials and Methods 2.5). In the current research no specific patient cohort stratification or selection was applied. All patients underwent planned colorectal or stomach tumour resection at Kyiv Region Cancer Center from October 2021 – January 2022.

One of the main challenges we sought to address is delivering a robust procedure for sample preparation. This would enable a reliable and consistent method of cell culture establishment, which is crucial for the biological evaluation of potential new chemical entities (**Figure 1a**).

**Figure 1.**
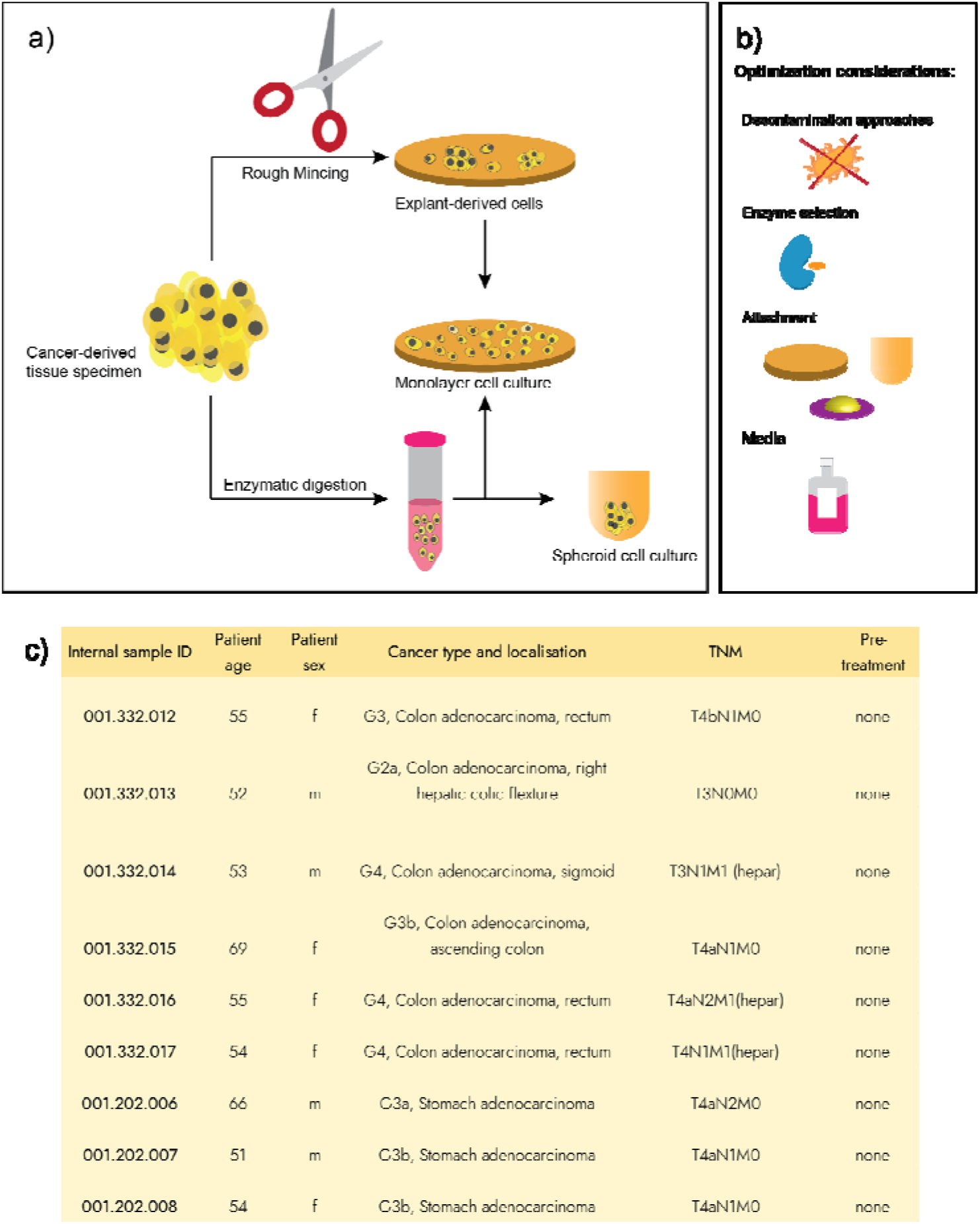
**a)** Generalized description of the cell isolation and primary cell culture establishment used in this work. **b)** Optimization approaches used in this work, **c)** generalized description of the samples, used in the current manuscript.

To establish a BioBank, viable cells with modest to high proliferation potential, an effective means of purification, and recovery from cryo-storage are required. There are many challenges in developing a viable method: these include cell isolation, removal of potential bacterial and fungal contamination, cell viability and yield, removal of necrotic apoptotic cells and reconstitution of cells from cryo-storage.

Decontamination of gastrointestinal tissues is one of the main challenges during the culture establishment. The superficial epithelial layer of the colon carries a diverse microflora population, which, if present, can compromise cell culture establishment.

Initially, we investigated the use of various alcohols for bacterial decontamination, which resulted in a significantly reduced cell yield and viability. We then evaluated several antibiotics in parallel, including a commercially available penicillin/streptomycin mix, Primocin^®^, Normocur^®^, Fungin^®^ (all – Invitrogen), and Amphotericin B.

Despite Penicillin/Streptomycin mixture being the most common antibacterial agent, in our hands it did not show sufficient reduction in bacterial contamination. For example, whilst preparing sample **001.332.012**, we experience overwhelming contamination. Thus, it was decided to evaluate Primocin^®^ and Normocur^®^ mixture. Encouragingly after a third digestion and seeding cycle. There was no evidence of bacterial growth during 27 days of culturing (4 passages).

A fungal contamination assessment was made by visual inspection. Initially, Amphotericin B was used to control fungal growth. However, subsequently Fungin^®^ was more effective. Therefore, we proceeded with Primocin^®^/Normocur^®^ and either Fungin^®^ or Amphotericin B for all tissue extractions to control microbial growth. In the light of reports (38) of a decrease in cell viability in the presence of Primocin^®^/Normocur^®,^ a modified procedure was developed. Two passages with Primocin^®^/Normocur^®^ and either Fungin^®^ or Amphotericin B, and a subsequent passage with Penicillin/streptomycin mixture treatment. This resulted in superior quality cultures.

The yield of isolated cells increased as the tissue samples were homogenized. Increase of total surface area in combination with softening of the tissue increases accessibility of the fragments to digestion enzymes. The enzymes Trypsin-EDTA, Dispase II, Hyaluronidase, and Collagenase Type IV were evaluated for their ability to digest the tissue. Collagenase Type IV (STEMCELL Technologies) in combination with Hyaluronidase (Sigma) had a similar effect to Dispase II. Both mixtures required the same time of exposure and temperature conditions for the complete digestion of the sample. Agitation was achieved either by shaking, vortex application, or pipetting of the sample in the digestion solution with a Pasteur pipette and were all equally effective during cell isolation.

The digestion of fibrotic tissues **001.332.015** and **001.202.007** proved challenging. It was not possible to produce specimen sizes of 1-2 mm in an acceptable period. Furthermore, for sample **001.332.015 a** lipid droplet phase separated from the overall mixture which hampered cell identification.

Removal of necrotic and apoptotic cells by visual evaluation of intracellular granulation, residual or emergent is a crucial step during culture maintenance. Since any of those cells remaining could negatively affect subsequent cell growth.

Certain cell types, including epithelial cells did not display efficient and rapid attachment to tissue culture plastic, including viable Trypan Blue negative cells. Some samples, *e*.*g*., **001.332.016** and **001.202.006** had more than 80% of cancer cells in suspension.

The explant-based technique is the major alternative to the suspension-based seeding of 2D monolayer after tissue digestion. We hypothesized that the absence of the enzymatic cleavage of the extracellular matrix proteins would improve attachment efficiency of the epithelial cells. Main variables for the optimization of the explant adaptation include the size of the final fragments, their density, and the time of exposure. At the same time, explant cultivation requires closer monitoring of the suspension and growth condition, due to the decomposition of the dead cells and non-cellular elements of the extracellular matrix.

For cultures **001.332.015, 001.332.016, 001.332.017, 001.202.007**, and **001.202.008** explant-based procedures resulted in higher attachment rates, and more natural cell physiology (**Table S5**) (**Figure S3**). Additionally, for **001.202.007** and **001.202.008** explant-derived cells formed 50-100 μm clusters, which in the case of **001.202.008** were expandable (**Figure 1b**).

We have also attempted the production of primary cell culture from cryopreserved cultures, which is a critical step in establishing a viable biobank. Tissues **001.212.007** and **001.212.008** were cryopreserved in standard conditions, DMEM-F12 80%, FBS 10%, DMSO 10% at -80°C, and stored for three months. Upon thawing the tissue was processed, following the digestion-based protocol. Cell attachment was found to be at 65%. Unfortunately, after two weeks in culture, attachment decreased to 50%, the cells in suspension appeared necrotic, and the culture was no longer viable.

### 3.2. 2D-cell culture conditions optimization

We found that each cell type culture required specific optimization. For cancer-associated fibroblast cultures **001.332.015f** and **001.202.008f**, tissue-culture coated plastic plates worked well, whereas epithelial phenotype cells did not. Typically, more than 70% of cells floated in the culture medium after initial seeding. Surprisingly, Trypan Blue staining did not show any difference in viability between the attached cells and cells floating in the medium -even after the week of cultivation. Furthermore, there was no evidence of significant granulation, deformation, or any other senescence-related phenotypic effects in floating cells (**Figure S3**). We postulated that those cells adapted a low-attachment phenotype. Moreover, they portrayed defined cell-cell interactions by forming floating spheres.

Two native xeno-derived Extracellular matrices ECMs for cell attachment were evaluated, Matrigel^®^ (Corning) and BME^®^ (Cultrex). Of the two, Matrigel^®^ is softer, which in our hands results in inconsistent plate coating and a lack of robustness to the mechanical stresses that occurs during culture maintenance. In contrast BME^®^ is more rigid and therefore less susceptible to mechanical stress during cultivation. Thus, we preferred BME^®^. A serial dilution experiments using the cell attachment as comparator, determined a 1:40 dilution of BME^®^ in the DMEM to be optimal for the thin layer coating, prior to seedng with cells (**Figure S4**).

Although fibrotic tissues samples **001.332.015** and **001.202.008** contain significant numbers of CAFs, treatment with explant-based techniques and subsequent differential detachment maximized the yield of cancer cells in culture. Nevertheless, after two weeks of growth they suffered from unacceptable levels of CAF growth.

Unlike the immortalized cell lines, primary cells usually do not reach confluent monolayer rapidly. However, BME^®^ monolayer slowly dissolves in media and detaches from the surface over 2 weeks. Thus, cell passage was executed in three cases:

a. if local confluence was higher than 90% in a region of more than 0.5 mm,
b. if cell detachment was stable for more than 1 week and thus culture required densification,
c. if matrix was damaged and required passaging.

A Simple wash with 50 μL EDTA with PBS at 4°C was enough to detach the cells. The detachment was validated microscopically.

If after two-weeks cells had not reached local confluence, cells were cryopreserved “as is”.

Cell viability was controlled throughout the procedure using two approaches. One, interim control with Trypan Blue staining. Cells were dismissed if the viability after the isolation was lower than 85%. Final establishment QC included Propidium Iodide staining of cells and subsequent analysis using fluorescent imaging (see Supporting Information for showcasing examples) (see **SI, Figure S9**).

Table at **Figure 2(f)** represents the time course of cells in culture and the maximum passage of ocassion (**a**) used for these cells. For some cultures, the passage counting is subjective due to the application of passages on occasions (**b**) and (**c**).

**Figure 2.**
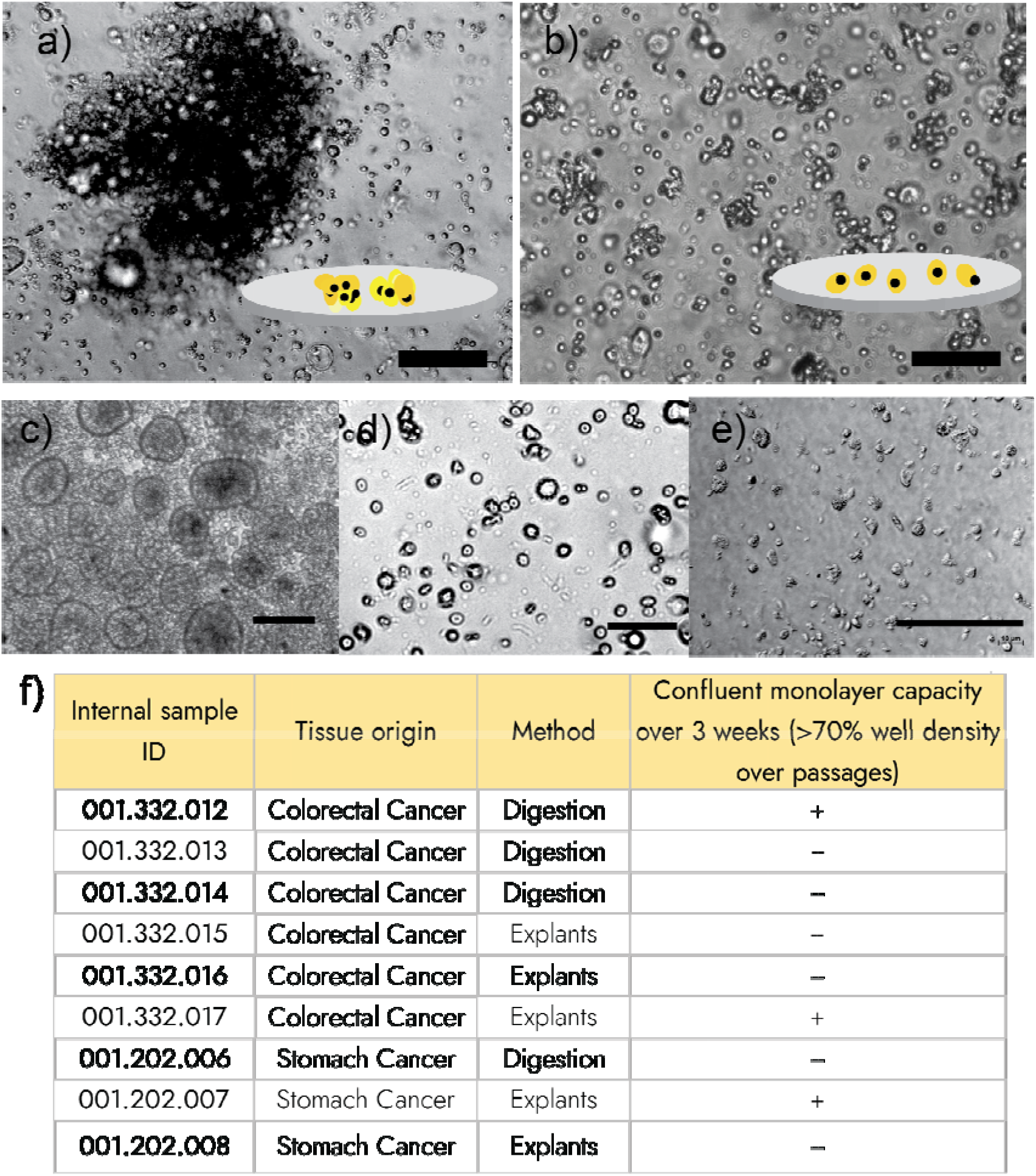
**(a)** Explant of the culture **001.202.007** in culture, cartoon represents the explant derivation; **(b)** Digestion-derived culture **001.202.007** after 3 days in culture, cartoon represents the digestion-based derivation; **(c)** Monolayer of the culture **001.202.007** at confluency, showing an abundance of organoid-like phenotypes; **(d)** Cells derived from culture **001.332.015** after 5 days in culture; **(e)** Cells derived from culture **001.332.017** after 4 days in culture. **(f)** Summary of cell expansion in attached monolayers. Scale bars on all images are at 50 μm.

The epithelial nature of the obtained 2D cell culture was shown on the example of culture **001.202.008** after 2 weeks after the attachment using EpCAM (epithelial phenotype marker) and CD44 staining, mesenchymal phenotype marker (see **SI**). Cells reproducibly showed higher expression of EpCAM, which confirms low mesenchymal phenotype of the formed culture (see **SI, Figure S8**).

### 3.3. Media optimization

Initially cultivation media optimization from serum-free medium formulation, using either RPMI, DMEM or DMEM/F12 was investigated. However, cultivation in serum-free media led to high levels of cell detachment. Addition of 10% FBS, a source of calcium and magnesium ions essential for effective cell attachment, to the cell culture media increased cell surface adhesion by up to 60% compared with cells cultured without FBS.

Several conditions were explored in previous in-house experiments with the aim of increasing the proliferation potential and decreasing cell death during cell culture. EGF, hydrocortisone and insulin, (as drivers of cell division, metabolism, and growth), along with transferrin (which inhibits free radical formation and protects cells from such damage) were added to the cell culture media. L-Glutamine was used as an essential precursor for new protein synthesis and one of the main members of the metabolic pathways inside the cell.

Hence, after different media composition trials, the optimal composition of media for the GIAC primary cell cultivation was: DMEM/F12 as a base, with the addition of L-glutamine, antibiotics, EGF, and B-27 supplement as a source of transferrin, insulin, and hydrocortisone with 10% FBS.

#### Cryogenic preservation

As a main component of the media for cell suspension freezing, PBS (80%) + FBS (10%) + DMSO (10%), DPBS (80%) + FBS (10%) + DMSO (10%), DMEM (90%) + DMSO (10%), FBS (90%) + DMSO (10%), and DMEM (90%) +FBS (10%) + DMSO (10%) for culture **001.332.012** were all investigated (see **Table S6**). Both PBS and DPBS with the addition of FBS 10% showed moderate results. The number of dead cells in the suspension after cryopreservation with these two buffers comprises 20%. Using DMEM with the addition of FBS (5:4 ratio) or FBS alone improved cell viability rate after the freezing cycle of the cells (95% and 96% for DMEM+FBS and pure FBS, respectively).

For freezing Mr. Frosty^®^ containers were used, which prolonged the freezing process over time at a maximum rate of 1°C/min. Freezing was accomplished in serval stages: First, samples (in vials) were cooled to 4°C for 30 minutes, then transferred to a -20°C freezer for 60 minutes and finally the temperature was reduced to -80°C overnight. For short-term storage we found liquid nitrogen to be more harmful to the cells.

Culture **001.332.012** was the most prolific of the cultures under investigation, it generated local confluent monolayers within 7-9 days. In contrast, other colorectal cell cultures, specifically **001.332.013, 001.332.014** and **001.332.015** did not display a similar growth profile and therefore were discarded after two weeks in culture (**Table S2**).

Culture **001.202.007** grew rapidly and developed into an organoid-like phenotype, reaching confluence inafter 7 days in culture (**Figure 2c**). By confluence here we mean the density of organoid spheres within the plate, where additional growth is no longer reachable.

The monolayer-based method resulted in low success rates (<25%). Therefore, it is not an appropriate general method

### 3.4. 3D-spheroid cell culture scalability

In contrast to 2D adherent monolayer cell growth, 3D-culture conditions require an environment that will increase cell-cell interactions. Two distinct approaches are known in this field: scaffold embedding (39) and low-attachment environment application (40). Matrigel scaffold embedding, an application of specialized media reported in (41), was investigated for the three cell cultures **001.332.012-014** and organoid-like structures were observed in all three samples. However, during subsequent passages of those cultures the organoids lost their viability due to the senescence (**Table S4**).

Classical spheroid formation was then evaluated using Ultra-Low Attachment (ULA) well-plates (CellCarrier Spheroid ULA 96-well microplates) and an overhanging drop procedure using the same media as in 2D culture establishment. The media was replenished to a half every second day.

Using this experimental set-up, non-adherent culture of **001.332.012** was initially evaluated. In contrast to previous experience with this culture this time it was not successful: doubling times in ULA conditions were estimated at 5 days. Furthermore, it did not form defined spheroids, and showed prominent peripheral growth over the monitoring period. The culture was expanded for three consecutive times, using this procedure. Short-term cultures in 2D format of **001.332.012, 001.332.015**, and **001.212.006**, in contrast, were expandable without attachment in 3D format (**Figure 3**). Cells were passaged every 6-9 days without significant loss of viability and cryopreserved at P3.

**Figure 3.**
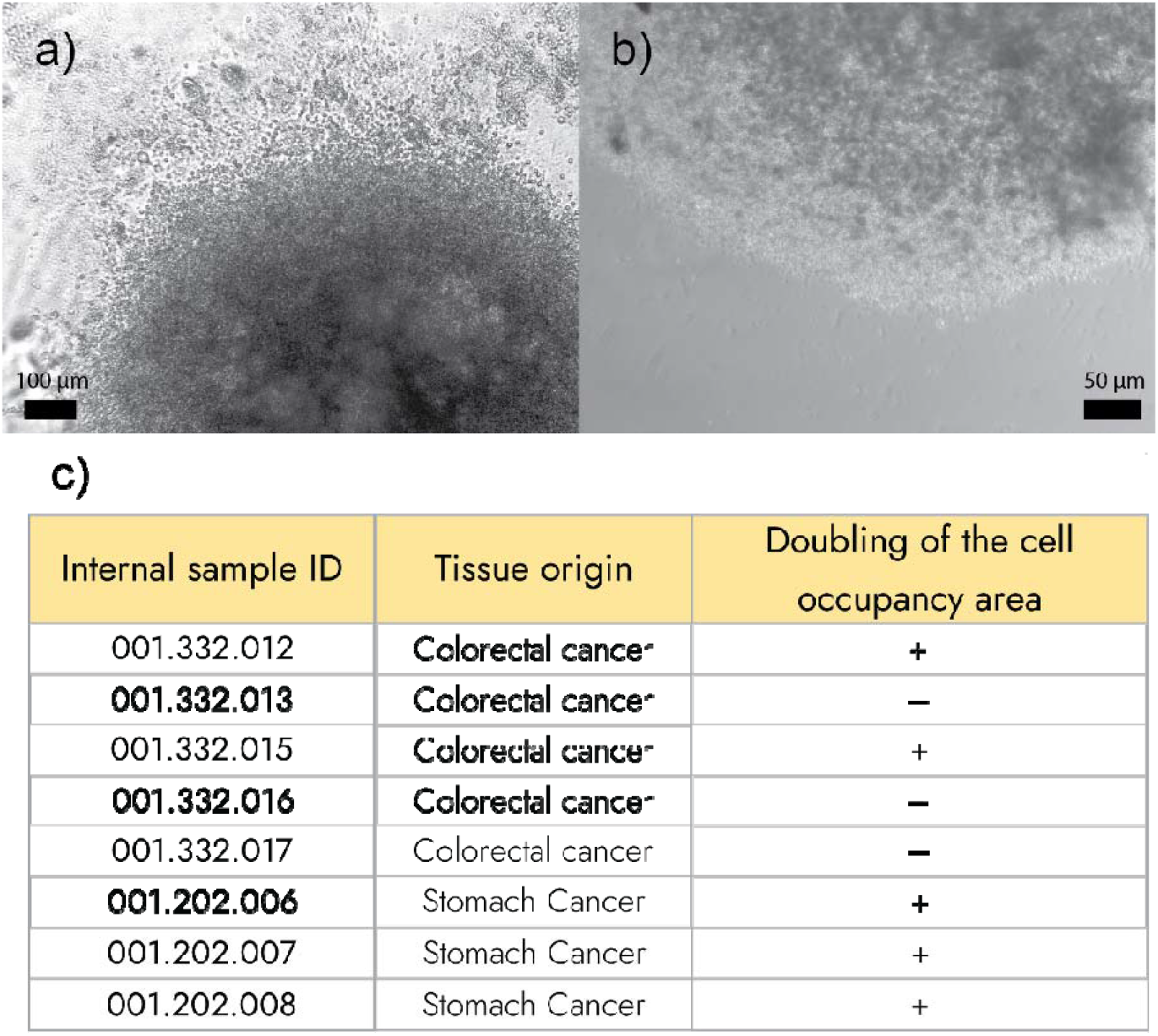
**(a)** Spheroids formed using the culture of **001.202.006** tissue; **(b)** and of the culture **001.332.015; (c)** Summary table showing growth potential of tested cells upon low-attachment phenotype stimulation.

While culture **001.332.012** formed ∼50 μm in size tumour spheres, cells **001.332.008** formed large millimeter-size homogeneous spheroids, which, could be broken down into fragments, and these further expanded. A similar observation was made for other cultures, described in **Figure 3c**.

In contrast to earlier observations, culture **001.332.008**, which had the highest level of fibroblast overgrowth in 2D, showed clear epithelial cells proliferation in the non-adherent 3D environment, and efficiently formed complete spheroids.

Generally, the increased proliferation of cell digestion in ULA conditions in contrast to 2D adherent monolayer conditions can be explained with increased content of cancer stem cells. Experiment with staining of ULA-spheroid with CD133 fluorescent antibody showed prevalence of CD133+ cells on ULA plates in contrast with regular BME^®^ -coated well-plates (**Figure S5**).

### 3.5. Cancer-Associated Fibroblasts isolation

CAFs are an important source of information on cancer biology. Nevertheless, their use in lead optimization and preclinical stages of cancer research is under-utilized.

Tissue stroma is the usual source of fibroblasts and the presence of >50% of stroma in the histological sample was a good predictive marker for successful CAF isolation in our hands (**Figure 4 a, b**).

**Figure 4.**
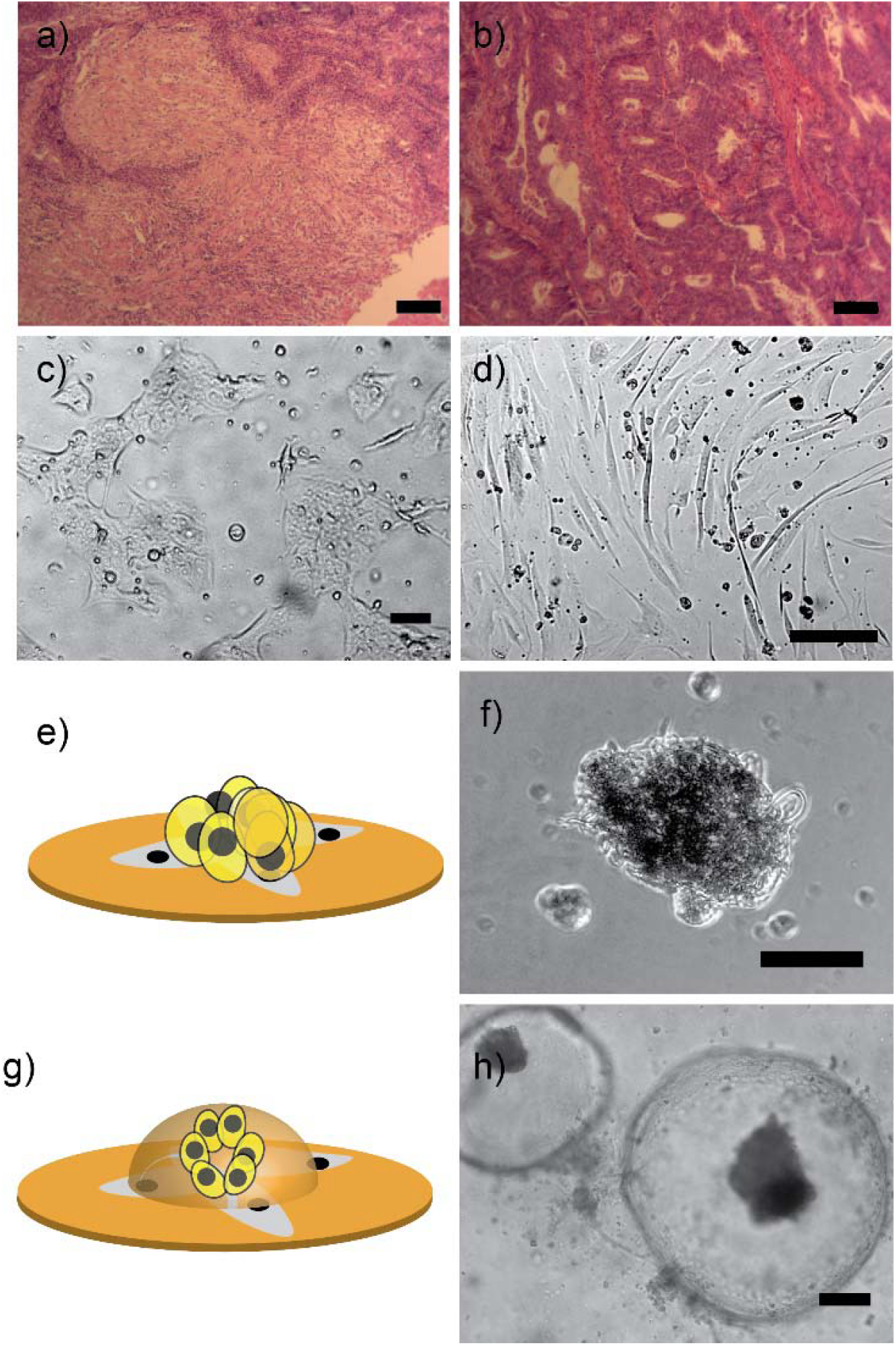
**(a)** Hematoxylin-eosin staining of an histological slice from specimen **001.212.008**, with stroma-rich regions (pinkish); **(b)** Histological slice of cancer cell rich (bordeux-colored) specimen **001.212.006**, with little or no stroma regions; (c) Image of the cancer-associated fibroblast colonies from culture **001.212.008; (d)** Image of the cancer-associated fibroblasts derived from the specimen **001.332.015; (e)** Schematic representation for epithelial cells spheroid grown on the top of cancer-associated fibroblasts layer; **(f)** defined epithelia cell spheroid, formed on the top of cancer-associated fibroblasts layer. Scale bar on all images are at 50 μm.

Initially sample **001.332.015** was used for evaluating optimization procedures and subsequent monitoring CAF behavior. This specimen was highly differentiated from other colorectal samples as judged by its density. The sample proved difficult to dissociate, hence as an alternative it was treated with the use of frequent (every 20 min) exchanges of Collagenase IV and Hyaluronidase solution. This resulted in 10 million viable cells used for plating.

On day two the formation of fibroblast colonies, mixed with colonies of epithelial cells were observed. After 2 weeks in culture, ∼30% of the overall cell population were fibroblasts within colonies. After treatment with gentle trypsinization (1 min), ∼80% of epithelial cells were removed. Subsequent 5 minute incubation with TrypLE Express resulted in complete detachment of fibroblasts from the bottom of the well-plate.

After the reseeding of the cells and overnight incubation, fibroblasts as individual cells (**Figure 4d**) were observed Those cells were successfully passaged 4 times, at which time a less definitive cytoskeleton structure and more granulation within the cells developed.

Interestingly, when epithelial cells were admixed to the fibroblast culture, they formed a spheroid-like structure on top of fibroblast layer. Those structures were stable within the passage and could be retained after passage on top of confluent layers of fibroblasts up to passage 4. A closer examination of those spheroids (**Figure 4f**) revealed the presence of ECM fibers.

CAFs were isolated from the highly stroma-rich tumour **001.332.008 (Figure 4d)**. However, those tended to remain within densely populated colonies, despite harsh digestion attempts between passages.

It should be noted that CAFs can be used in the co-culture with the cancer-derived spheroid culture on Matrigel. CAFs growth occur in parallel to the spheroid growth in this case, while spheroids adapting morphologically relevant phenotypes (**Figure 4g,h**).

## 4. Discussion

2D monolayer cancer cell cultures did not show the reliable proliferation rates for the cell culture growth on the scale of 9 patient-derived cells, and according to our results is not a model of choice for the creation of diversity-oriented cell arrays. Comparing digestion and explant-based growth, we can conclude that explants are more appropriate for cell-based assays using those specific primary cells and can be used for short-term experiments. Digestion-derived cells show moderate-to-low attachment characteristics. Explant-based cells attached to BME-coated well-plates successfully in most of the cases, and reached confluency in some examples. That can be attributed to the conservation of collagen matrix within conglomerates itself, and, perhaps, less harmful and faster processing. Some tissues, specifically, with 50% or more fibrotic tissue in histological slide cannot be digested within the 3 hours, thus, explants remain the only culture establishment choice.

Those, however, are complicated to monitor using modern imaging techniques, since the fluorescent staining is inhomogeneous and strongly depends on the explant preparation procedure. Tissue characteristics are important for those models as well, since fibrotic tumour specimens quickly become unusable in the cancer therapeutics screening, since fibroblasts overgrow tumour cells.

3D spheroid cell models, in contrary showed stable proliferation rates within our experimental set-ups. Those cell cultures showed cell occupancy doubling times of 7-14 days and could be passaged successfully. These data are in accordance to the previous findings (32,42,43) and this work can serve as a prove of the validity of those findings. Formation of spheres or large-size (100 μm or more) spheroids can both occur, depending on the tissue. That finding can be, perhaps, explained by cell-cell interaction protein expression and specific cell phenotypes. That analysis, however, is out of the scope of this work.

We used the homogeneous media for all experiments, which may simplify our attempts to establish the stable and cost-effective cell culture isolation procedure for the cell growth.

Cancer-associated fibroblasts can be isolated according to the procedures, specific for 2D monolayer cell cultures. Those quickly overgrow epithelial tumour cells and become dominating within the monolayer. Curiously cancer cells were shown sometimes to adapt the spheroid morphology on the top of the fibroblast monolayers. Moreover, those can be combined with the BME-encapsulated cell culture to generate the morphologically relevant microtissue models, which hopefully may be useful in the future research.

## 5. Conclusions

This practical assesment of modern cell culturing techniques in application to GIAC, clearly demonstrated that 2D adherent monolayer is an inadequate method for use in the modeling of specific diseases. Despite monolayers being preferable for cell observation and assay design, this format cannot guarantee an efficient expansion of cancer cells. In contrast, for CAFs this method guarantees homogeneous expansion and purification during culturing.

The 3D organoid method resulted in the formation of the organoid-like structures during the initial stages in culture. However, these quickly entered senescence after passaging.

In contrast, the 3D spheroid procedure, demonstrated the best results both in terms of avoidance of early cell senescence and estimated doubling time. This procedure helps undifferentiated proliferating phenotypes to exist longer than in 2D adherent monolayer set up.

Primary cells may not always form a defined spheroid, instead remaining in 70-100 μm aggregates. However, both phenotypes have the potential to expand and be used in living-cell biobanking.

## Supplementary Materials

Results and experimental procedures that augment this manuscript can be found at xxx/bmc.com as Supporting Information.

## Author Contributions

Conceptualization, A.I.H., D.M.V., I.T; methodology, I.T., O.M.; validation, A.D., O.A.K., D.O.S., O.M.; investigation, I.T., O.M, O.A.K., A.D.; resources, S.O.V., I.S.G., O.A.R.; writing—original draft preparation, A.I.H.; writing—review and editing, D.M.V., O.M., S.V.R., D.B.J.; supervision, A.I.H.; project administration, O.A.R., I.S.G., I.T., A.I.H. All authors have read and agreed to the published version of the manuscript.

## Institutional Review Board Statement

The study was conducted in accordance with the Declaration of Helsinki and approved by the Ethics Committee of Kyiv Regional Oncology Center (January 8^th^ 2021).

## Informed Consent Statement

Informed consent, approved by an Ethical Committee was obtained from all subjects involved in the study.

## Data Availability Statement

The data of this study are available from the corresponding author upon request.

## Acknowledgments

We thank the Biopharma Plasma LLC CEO and Founder of Preci LLC Konstiantyn Yefymenko for the generous support during this project. His valuable insights and discussions were crucial for strategic planning of this project. We also acknowledge the valuable input from Biopharma Plasma CTO Serhii Kovalchuk, whose precise advises allowed to progress in this project. Research was conducted at Preci LLC. Authors also would like to thank to the Cellesce Ltd. for the motivation to conduct this work. Ukrainian scientists thank all brave Ukrainian soldiers who gave a possibility to finish their research.

