## Supplementary Data for "Recent developments in the production of 2D- and 3D colon and stomach adenocarcinomas primary cell models"

**For the manuscript**

**Table S1. Comprehensive summary of the biospecimen characteristics, used in the current study.**

| Internal sample ID | Tumor mass | Detailed tissue description | Cell yield |
|---|---|---|---|
| 001.332.012 | 5g | Yellow soft tissue with a heap of blood vessels, fat, and blood. Without necrosis | 6.5 Mio |
| 001.332.013 | 4g | Stiff cartilage-like orange tissue with a bit of blood and blood vessels. Without fat or necrosis regions. | 5.2 Mio |
| 001.332.014 | 3g | Reddish soft tissue with a heap of blood vessels, necrosis regions, and blood. Without fat | 1.5 Mio |
| 001.332.015 | 5g | Yellow stiff as cartilage tissue with a heap of blood vessels, fat, and blood. Without necrosis | 10 Mio |
| 001.332.016 | 4g | Orange reddish soft tissue with blood vessels and fat. Without necrosis. | 4.5 Mio |
| 001.332.017 | 4g | Yellow reddish tissue without blood vessels, blood, necrosis, or fat | 4 Mio |
| 001.202.006 | 3g | Orange reddish soft tissue with blood vessels and blood. Without necrosis and fat | 4.3 Mio |
| 001.202.007 | 2g | Yellow reddish soft tissue with blood vessels, necrosis, and fat. | 3.1 Mio |
| 001.202.008 | 6g | Reddish stiff cartilage-like tissue with blood and necrosis. Without fat. | 25 Mio |

**Table S2. Essential cell culture information (2D adherent monolayer cells).**

| Internal sample ID | Establishment method | Attachment at P0 | Confluent monolayer capacity over 3 weeks (>70% well density over passages) |
|---|---|---|---|
| 001.332.012 | Collagenase IV and Hyaluronidase | 95% | + |
| 001.332.013 | Collagenase IV and Hyaluronidase | 93% | – |
| 001.332.014 | Collagenase IV and Hyaluronidase | 85% | – |
| 001.332.015 | Explants culture | 97% | – |
| 001.332.016 | Collagenase IV and Hyaluronidase. Explants culture in parallel | 83% | – |
| 001.332.017 | Collagenase IV and Hyaluronidase. Explants culture in parallel | 92% | + |
| 001.202.006 | Dispase II | 85% | – |
| 001.202.007 | Collagenase IV and Hyaluronidase. Explants culture in parallel | 88% | + |
| 001.202.008 | Collagenase IV and Hyaluronidase. Explants culture in parallel | 96% | – |

**Table S3. Essential cell culture information (ULA conditions).**

| Internal sample ID | Establishment method | Proliferation capacity (>70% well density over passages) |
|---|---|---|
| 001.332.012 | Collagenase IV and Hyaluronidase | + |
| 001.332.013 | Collagenase IV and Hyaluronidase | – |
| 001.332.015 | Explants culture | + |
| 001.332.016 | Collagenase IV and Hyaluronidase. Explants culture in parallel | + |
| 001.332.017 | Collagenase IV and Hyaluronidase. Explants culture in parallel | + |
| 001.202.006 | Dispase II | – |
| 001.202.007 | Collagenase IV and Hyaluronidase. Explants culture in parallel | + |
| 001.202.008 | Collagenase IV and Hyaluronidase. Explants culture in parallel | + |

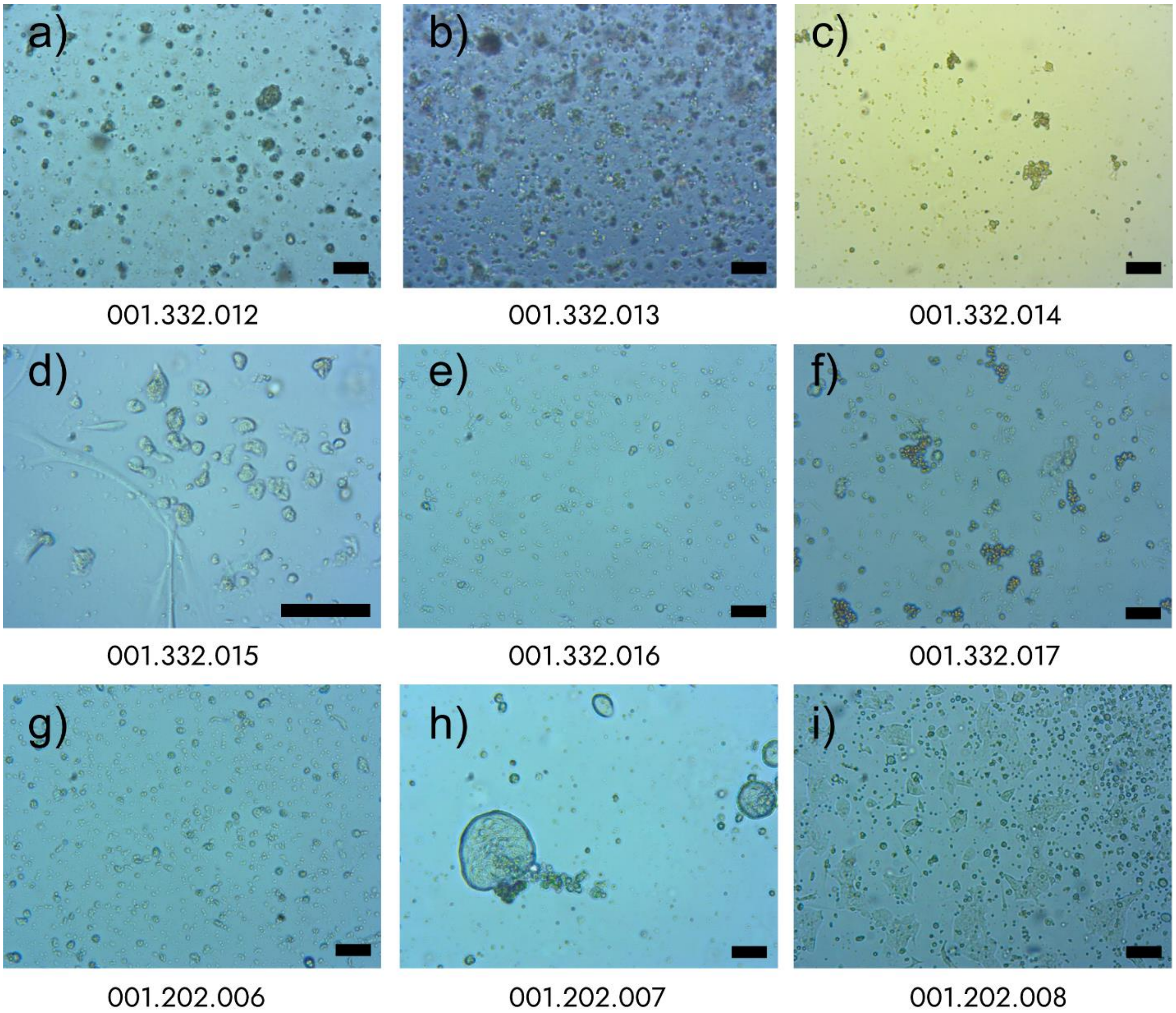


**Figure S1. Representative images of 2D cell cultures.** Colon adenocarcinoma (001.332) primary cell cultures grown as a monolayer: a) 012 at P1, 100x. b) 013 at P0, 100x. c) 014 at P2, 100x. d) 015 at P4, 400x. e) 016 at P1, 100x. f) 017 at P0, 100x. Stomach adenocarcinoma (001.202) primary 2D cell cultures: g) 006 at P1, 100x. h) 007 at P0, 100x. i) 008 at P0, 100x.

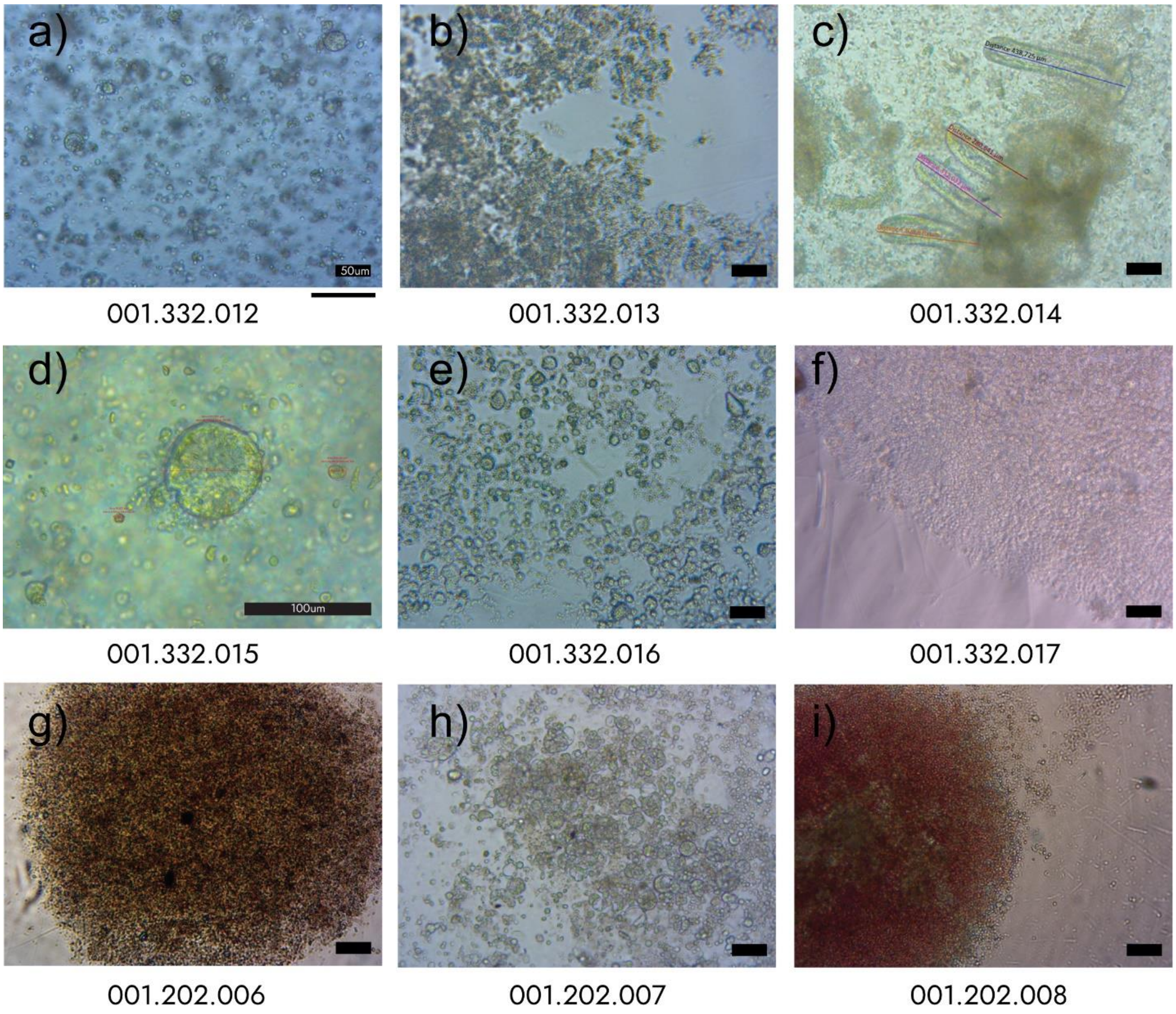


**Figure S2. Representative images of 3D cell cultures in domes/ULA conditions.** Colon (001.332) and stomach adenocarcinoma (001.202) primary cell cultures at P1-P4. Matrigel or BME were used as a scaffold for dome formation (a-d) and lead to the freezing of cell culture in current state. Ultra-low attachment plates-grown spheroids (e-f). Scale bar at 100 μm.

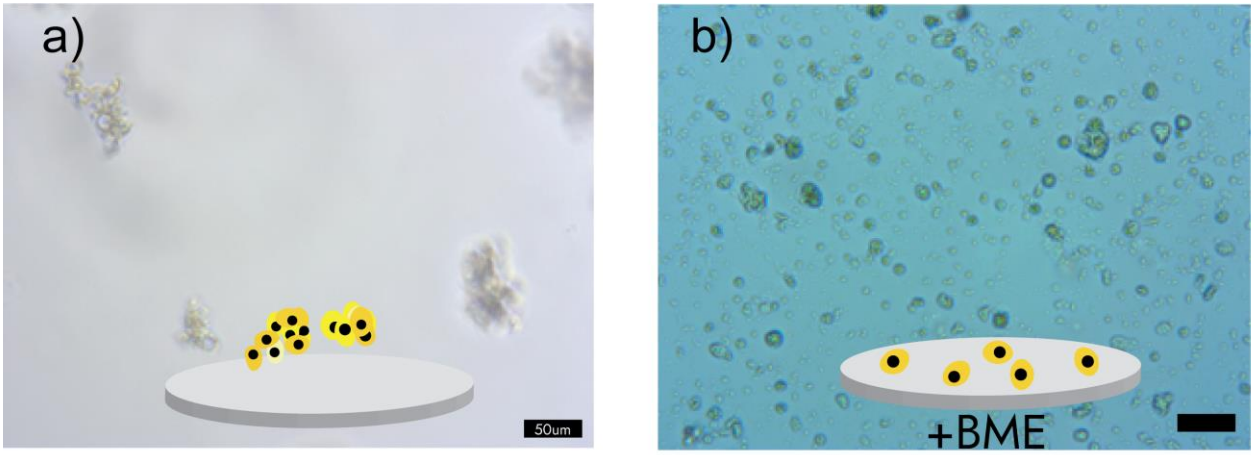


c)

| Internal sample ID | Attachment at P0 without pre-coating | Attachment at P1 after adding pre-coating step |
|---|---|---|
| 001.202.006 | 29% | 85% |
| 001.322.016 | 37% | 83% |

**Figure S3. Improved cell-surface attachment capacity after pre-coating usage.** a) Floating 001.202.006 cells in suspension without attachment to the surf ace. b) The same 001.202.006 cells on BME pre-coated surface. c) Summary table showing attachment rate with and without pre-coating.

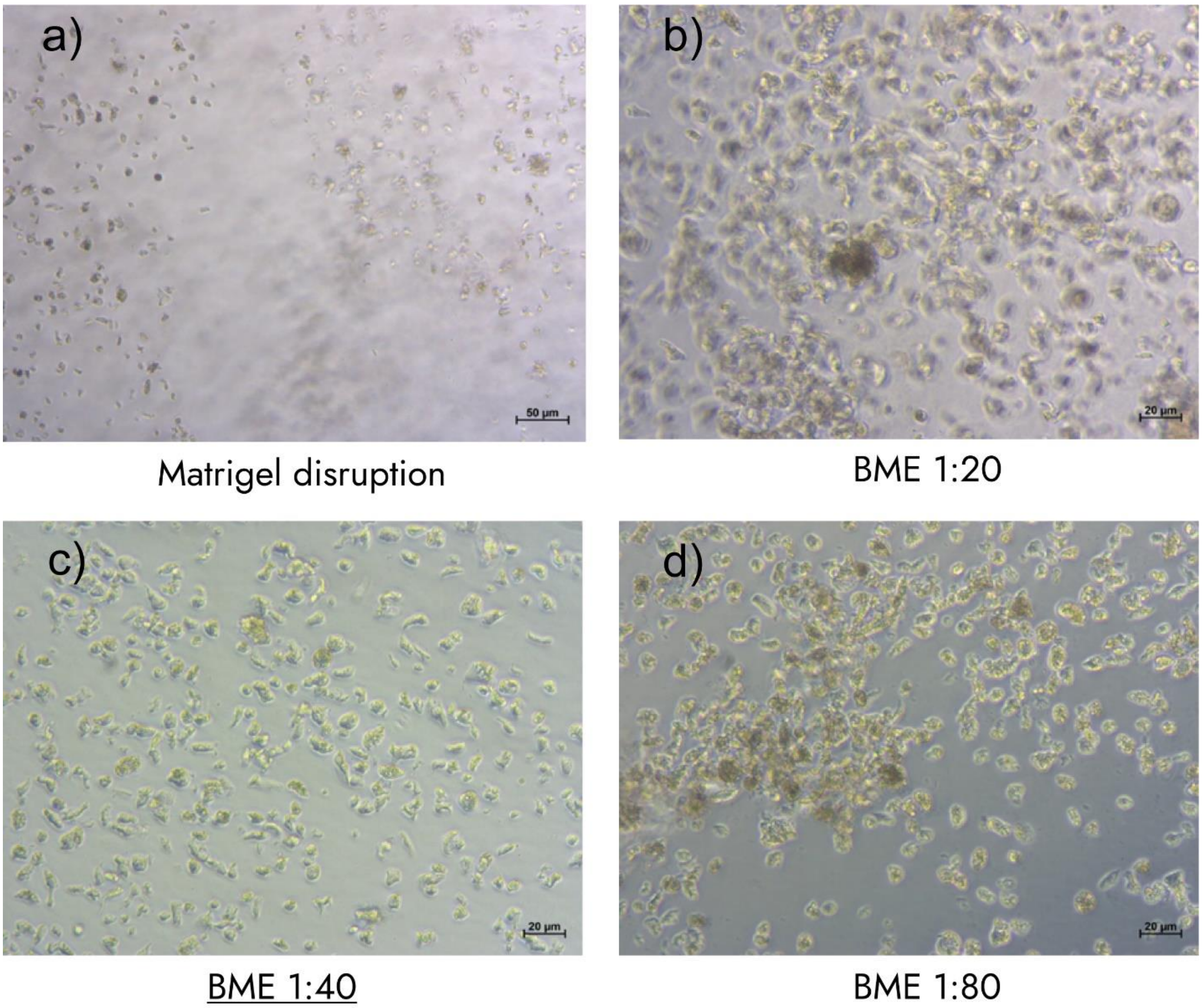


**Figure S4. Matrigel and BME pre-coating comparison.** a) Stable Matrigel coating (left part of an image), and disruption of Matrigel clump with the cells on it (right part of the image). b) BME 1:20 coating where the coated layer is too thick which leads to problems with focusing for monolayer. c) BME 1:40 coating with excellent cell distribution under the coating layer. d) BME 1:80 coating where clumps of BPE with the cells is floating on the medium.

**Table S4. Essential cell culture information (Matrigel/BME domes conditions).**

| Internal sample ID | Establishment method | Cells/conglomerates at P0 | Number of passages | Cell yield at Pmax |
|---|---|---|---|---|
| 001.332.012 | Collagenase IV and Hyaluronidase | 6,5 Mio/ 2,6K | 3 | Stopped due to conglomerates instability through passaging |
| 001.332.013 | TrypLE-EDTA | 5 Mio/12K | 4 | Stopped due to conglomerates instability through passaging |
| 001.332.014 | Collagenase IV and Hyaluronidase | 1.4 Mio/- | 2 | Stopped due to conglomerates instability through passaging |

**Table S5. Summary table showing attachment rate after enzymatic digestion and explant-based primary cell culture isolation.** Measurements were taken upon 48h plating period.

| Internal sample ID | Attachment after enzymatic digestion | Attachment after explants cultivation |
|---|---|---|
| 001.332.015 | 84% | 97% |
| 001.322.017 | 83% | 92% |
| 001.202.007 | 81% | 88% |
| 001.202.008 | 89% | 96% |

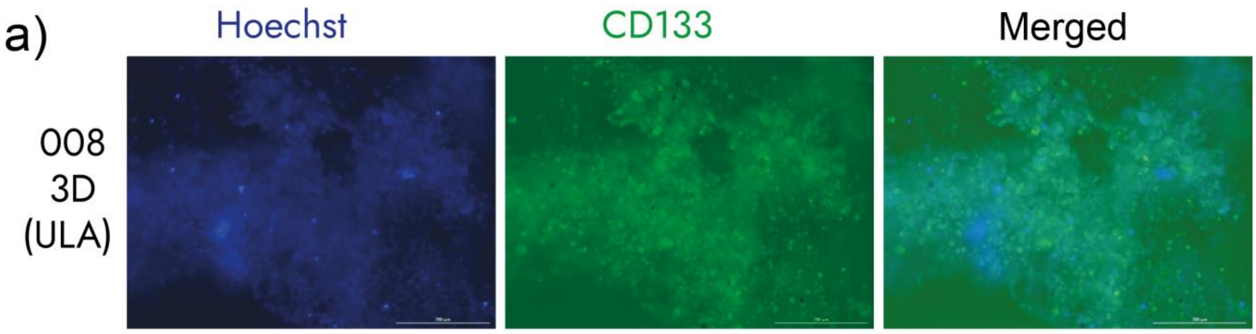


**Figure S5. CD133 expression abundance in ULA plated cells.** Following the gentle thawing procedure, 001.202.008 cells were examined for CD133 (stem cell marker) for rather low-attachment phenotype examination. (a) CD133 expression signal (green on the upper shots) was detected via immunofluorescence in 001.202.008 cells grown either in ULA or 2d monolayer settings for 5 days. Hoechst was used for cell visualization.

### Cell viability from frozen tissue

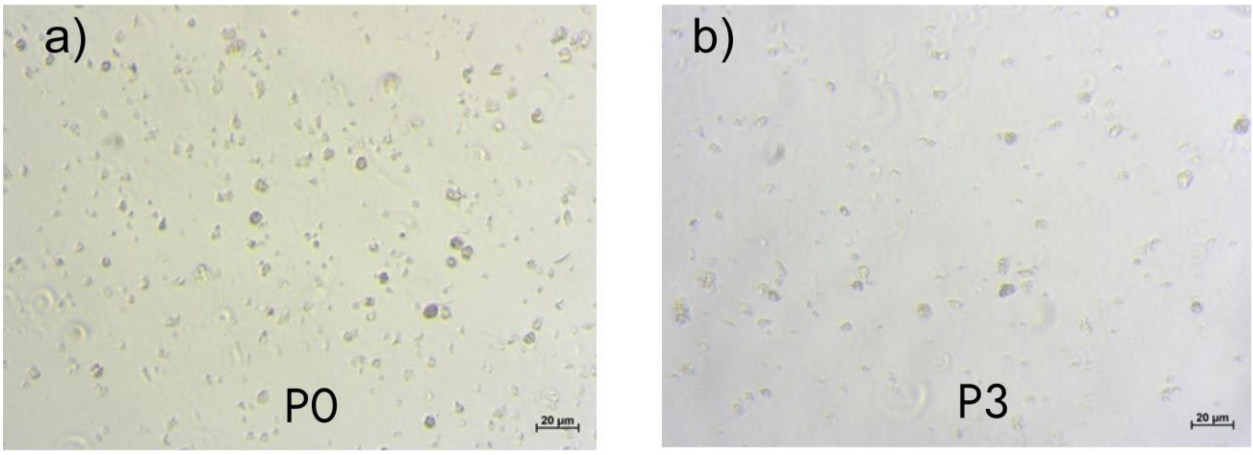


c)

| Internal sample ID | Attachment at P0 after frozen tissue digestion | Attachment at P3 after frozen tissue digestion | Trypan positive cells after digestion | Trypan positive at P3 |
|---|---|---|---|---|
| 001.202.007 | 63% | 36% | 17% | 59% |
| 001.202.008 | 71% | 58% | 19% | 41% |

**Figure S6. Viability of cells, extracted from frozen tissue samples.** a) Summary table showing cell viability. b) 001.202.007 cells at P0. c) 001.202.007 cells at P3.

### Freezing optimization

| Freezing medium | Viable cells after thawing |
|---|---|
| PBS:FBS:DMSO (8:1:1) | 79% |
| DPBS:FBS:DMSO (8:1:1) | 81% |
| DMEM:FBS:DMSO (8:1:1) | 93% |
| DMEM:FBS:DMSO (5:4:1) | 95% |
| DMEM:DMSO( 9:1) | 96% |
| FBS:DMSO(9:1) | 95% |

**Table S6. Summary table showing relation of freezing medium and cell viability after thawing.**

### Differential splitting of fibroblasts from epithelial cells

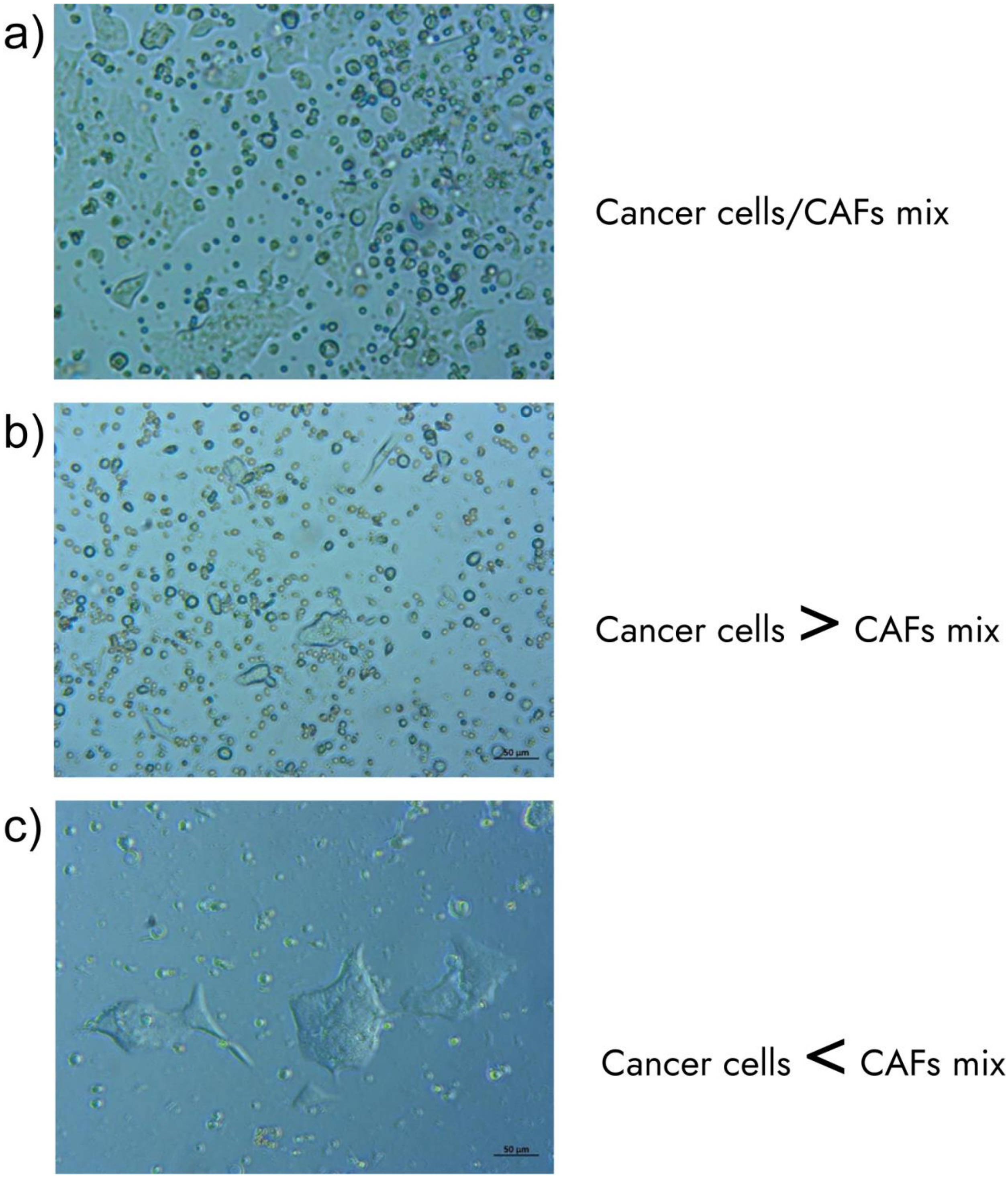


**Figure S7. Differential fibroblasts and epithelial cells splitting (001.202.008 culture).** Several techniques of detachment and centrifugation were tested to ensure safe separation of malignant and stromal cells within GIAC cultures. a) A mixture of fibroblasts and epithelial cells. b) Mostly epithelial cells after differential splitting. c) More clear CAF culture consisting of fibrotic conglomerates firmly attached to the surface.

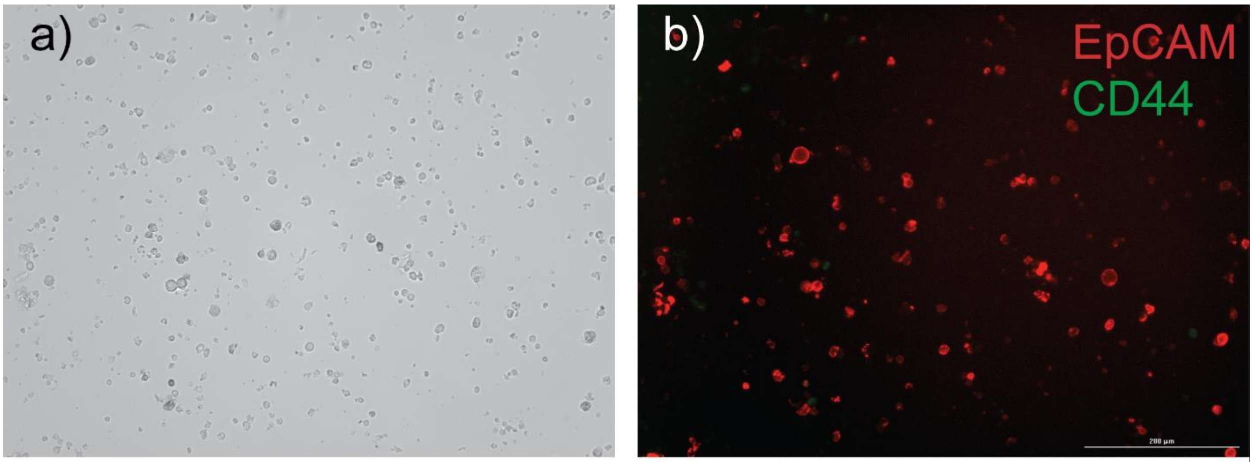


c)

| 001.202.008 | Avarage total cell number | Average number of positive cells | |
|---|---|---|---|
| | | EpCAM+ | CD44+ |
| Absolute values | 190 | 59 | 7 |
| Percentage | 100% | 31.05% | 3.68% |

**Figure S8. EpCAM and CD44 expression abundance in 2D plated cells.** Following the gentle thawing procedure, 001.202.008 cells were examined for CD44 (mesenchymal marker) and EpCAM (epithelial marker). (a) Brightfield image of cells grown in 2D. (b) EpCAM (red) and CD44 expression signals (green on the upper left shot) were detected via immunofluorescence in 001.202.008 cells grown on monolayer settings for 5 days. (c) Representative calculation from 3 independent experiments presented in a table.

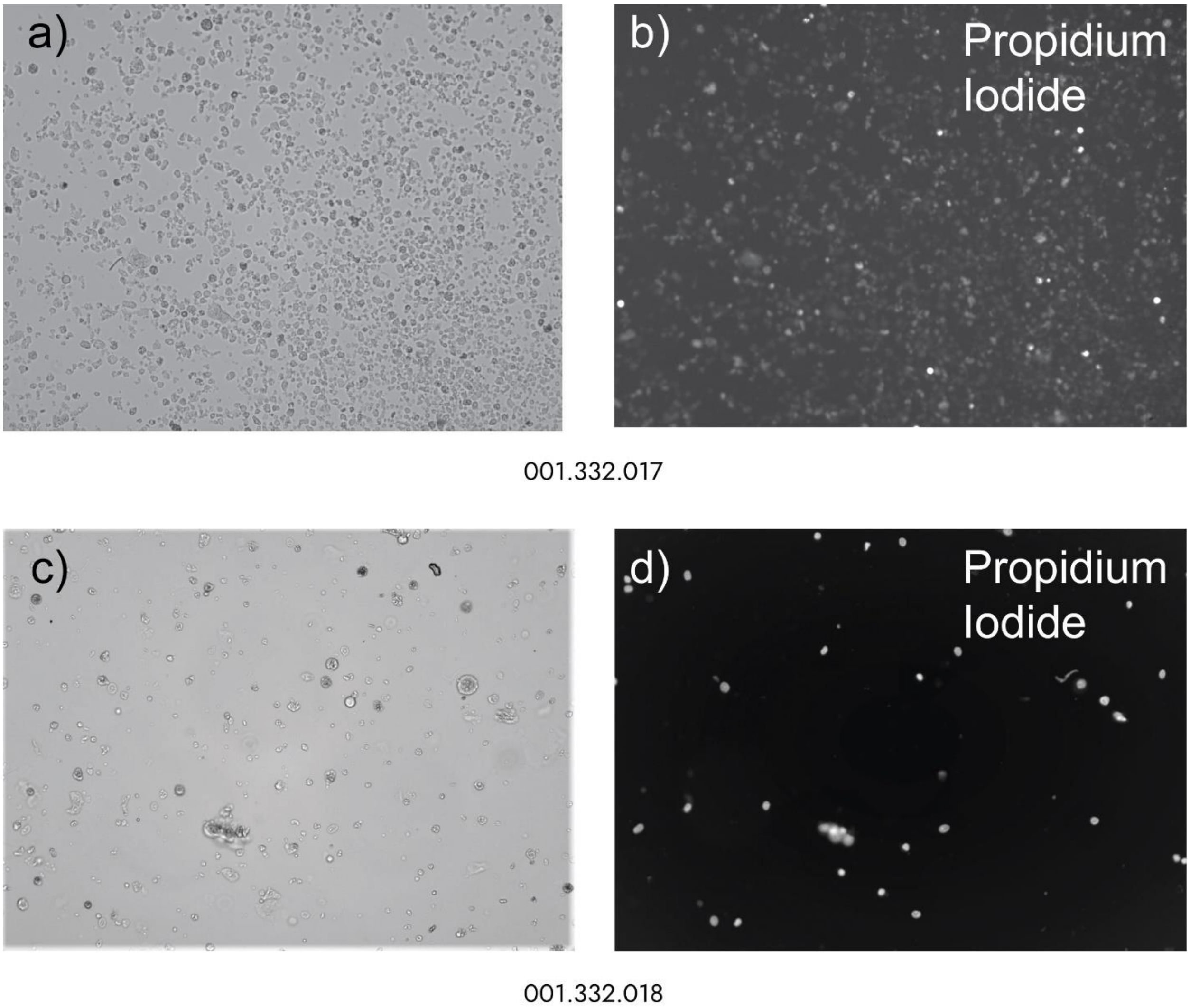


**Figure S9. Propidium Iodide staining.** Representative images of colorectal samples stained with Propidium Iodide (PI) (10mg/ml) by direct well addition. Subsequent fluorescent imaging conducted at 100x using Cytation Reader machine. Similar approach was routinely used for all isolated cells at different timepoints for dynamic cell viability testing. (a-b) 001.332.017 sample PI imaging after 10min of Ethanol-DMSO treatment. (c-d) 001.332.018 new sample viability assessment using PI staining after thawing.
